# LIS1 is critical for axon integrity in adult mice

**DOI:** 10.1101/2025.10.20.683562

**Authors:** Samaneh Matoo, Anne M. Ventrone, Shreena Patel, Jack K. Otterson, Sean Noonan, Noah Leever, Timothy J. Hines, Ashley L. Kalinski, Deanna S. Smith

## Abstract

Mutations in human LIS1 cause lissencephaly, a severe developmental brain malformation. Although most studies focus on development, LIS1 is also expressed in adult mouse tissues. We previously induced LIS1 knockout (iKO) in adult mice using a Cre-Lox approach with an actin promoter driving CreERT2 expression. This proved to be rapidly lethal, with evidence pointing toward nervous system dysfunction. CreERT2 activity was observed in astrocytes, brainstem and spinal motor neurons, and axons and Schwann cells in the sciatic and phrenic nerves, suggesting dysfunctional cardiorespiratory and motor circuits. However, it is unclear how LIS1 knockout in these different cell types contributes to the lethal phenotype. We now report that LIS1 depletion from astrocytes is not lethal to mice (male or female), although glial fibrillary protein (GFAP) expression is increased in all LIS1-depleted astrocytes. In contrast, LIS1 depletion from projection neurons causes motor deficits and rapid lethality in both males and females. This is accompanied by progressive, widespread axonal degeneration along the entire length of both motor and sensory axons. Interestingly, sensory neurons harvested from iKO mice initially extend axons in culture but soon develop axonal swellings and fragmentation, indicating axonal degeneration. LIS1 is a prominent regulator of cytoplasmic dynein 1 (dynein, hereafter), a microtubule motor whose disruption can cause both cortical malformations and later-onset neurodegenerative diseases, such as Charcot-Marie-Tooth disease. Our results raise the possibility that LIS1 depletion, through disruption of dynein function in mature axons, may lead to Wallerian-like axon degeneration without traumatic nerve injury.

**Significance Statement:** A healthy nervous system requires that proper brain wiring is maintained throughout the life of the animal. Connectivity often involves the long axons of projection neurons. Some axons drive cognition, others contribute to sensory and motor systems, while still others subserve vitally important cardiorespiratory processes. We show that LIS1, a protein linked to congenital brain abnormalities, also plays a crucial role in fully developed projection neurons in the adult mouse. LIS1 depletion from these cells causes severe axonal degeneration resembling the Wallerian degeneration that occurs in response to nerve injury. Because LIS1 regulates dynein, and because defective dynein can cause neurodegenerative disorders in humans, our study suggests that drugs targeting Wallerian degeneration may have therapeutic potential for dynein-related diseases.

## Introduction

*LIS1* haploinsufficiency causes lissencephaly, a developmental disorder associated with cognitive/motor impairments and short life expectancy (Di Donato et al., 2017; Brock et al., 2021). Homozygous disruption of *LIS1* in developing mice is peri-implantation lethal (Hirotsune et al., 1998). Loss of LIS1 protein interferes with the normal proliferation and migration of neural precursors, disrupts nuclear envelope breakdown and alters mitotic spindle dynamics (Gambello et al., 2003; McManus et al., 2004; Wynshaw-Boris, 2007; Hebbar et al., 2008; Reiner and Sapir, 2013; Moon et al., 2014). Heterozygotes do not exhibit the severe phenotypes associated with lissencephaly in humans, possibly due to size differences (Wynshaw-Boris et al., 2010).

LIS1 binds to and regulates dynein, a microtubule (MT) motor (Faulkner et al., 2000; Liu et al., 2000; Smith et al., 2000), and mutations in dynein heavy chain (*DYNC1H1*) can also cause lissencephaly (Marzo et al., 2019; Moller et al., 2024).The proportion of active dynein motors in a cell is controlled by LIS1 (Baumbach et al., 2017; Gutierrez et al., 2017; Elshenawy et al., 2020; Markus et al., 2020; Marzo et al., 2020; Xiang and Qiu, 2020; Singh et al., 2024). Dynein exists in a closed, autoinhibited form (the *phi* conformation) or an open activatable form (Zhang et al., 2017). Only the open form associates with the dynactin complex and cargo adaptors, interactions that are required for processive movement. LIS1 prevents dynein from adopting the autoinhibited form, promotes transition to the open form, and promotes the recruitment of multiple motors per dynactin complex, increasing motor speeds and run lengths (Htet et al., 2020; Markus et al., 2020; Rao et al., 2025). LIS1 interacts with dynactin’s p150 subunit, stabilizing dynein-dynactin complexes (Singh et al., 2024). These studies indicate that the proportion of dynein motors that are “activatable”, as well as their speeds and run lengths, can be influenced by LIS1 concentration.

We showed that LIS1 also has post-developmental roles in mice, using an actin promoter to drive a tamoxifen (TX)-inducible CreERT2 recombinase globally in adult mice with heterozygous or homozygous floxed LIS1 alleles (Hines et al., 2018). Heterozygous iKO mice (TX-*Act-LIS1^fl/wt^*) had no obvious phenotypes and survived long term, despite widespread CreERT2 activity by two weeks, indicating that loss of one LIS1 allele is tolerated in adult mice. In contrast, homozygous deletion of LIS1 (TX-*Act-LIS1^fl/fl^*) resulted in the rapid onset of severe neurological phenotypes, and mice died soon after TX administration. At peak phenotype severity, a CreERT2 activity reporter showed prominent recombinase activity in the heart but only sparse activity in the nervous system and other tissues. A Myh6 promoter was used to drive CreERT2 in cardiomyocytes, but no phenotypes appeared, even long term.

Within the nervous system, CreERT2 activity was predominantly observed in astrocytes and some neurons in the brainstem, hindbrain and spinal cord. We posited that defective cardiorespiratory and motor circuits might underlie the TX-*Act-LIS1^fl/fl^* phenotypes. Because LIS1 regulates key dynein-dependent processes in neurons, including long-distance cargo transport within axons (Sasaki et al., 2000; Yamada et al., 2008; Pandey and Smith, 2011; Yi et al., 2011; Pandey et al., 2022; Fellows et al., 2024) and the development and function of the axon initial segment (Kuijpers et al., 2016; Klinman et al., 2017; Torii et al., 2020), Some of the phenotypes were probably due to LIS1 depletion in neurons. However, more astrocytes than neurons showed evidence of CreERT2 activation, raising the possibility that LIS1 depletion in astrocytes contributed to the iKO phenotypes. To determine the cell-type-specific consequences of LIS1 depletion we have now used an Aldh1l1 promoter to drive CreERT2 expression in astrocytes (TX-*Aldh1l1-LIS1^fl/fl^*mice) and a Thy1 promoter to drive CreERT2 expression in projection neurons (TX-*Thy1-LIS1^fl/fl^*). The results unequivocally demonstrate a vital role for LIS1 in projection neurons and indicate that LIS1 depletion in these cells caused the lethality in TX-*Act-LIS1^fl/fl^*. Although LIS1 depletion in adult astrocytes did not produce obvious neurological symptoms or lethality, cellular changes suggest important roles for LIS1 in these cells.

## Materials and Methods

### Animal Use

Animals were bred and housed at the University of South Carolina Animal Resources Facility. The following strains purchased from Jackson Laboratory were used to generate two iKO strains:

**1*29S-Pafah1b1^tm2Awb^/*J:** (Jackson Laboratory 008002, RRID: IMSR_JAX:008002). Mice have loxP sites flanking exons 3–6 of the *Lis1* gene. This results in LIS1 depletion when mice are exposed to Tamoxifen (TX).

***B6; FVB-Tg (Aldh1l1-cre/ERT2)1Khakh/J:*** (Jackson Laboratory 029655, RRID: IMSR_JAX:029655. Mice express a CreERT2 fusion protein in astrocytes. CreERT2 becomes activated and moves to the nucleus upon TX administration.

***Tg (Thy1-cre/ERT2-EYFP) HGfng/PyngJ:*** (Jackson Laboratory 012708, RRID: IMSR_JAX:012708). Mice, also known as SLICK-H, express CreERT2 and enhanced yellow fluorescent protein (EYFP), each under the control of two separate modified Thy1 promoters. All projection neurons express EYFP. TX activates the CreERT2 in the same neurons.

***B6. Gt(ROSA)26Sor^tm9(CAG-tdTomato)^ ^Hze^/J:*** (Jackson Laboratory 007909, RRID: IMSR_JAX:007909): Mice, also known as Ai9, serve as a CreERT2 recombinase activity reporter; CreERT2 activity removes a loxP-flanked STOP cassette that prevents transcription of the CAG promoter-driven tdTomato. CreERT2 activity induced by TX results in robust tdTomato fluorescence. Genotyping of all animals was conducted using primers and protocols provided by The Jackson Laboratory, with primers available upon request. Males and Female mice were used, with no differences observed in iKO phenotypes based on sex of the animals. The designation for experimental and control animals is shown in the Table 1 with the most commonly used controls being the no floxed LIS1 mice exposed to TX.

### Tamoxifen (TX) administration

Tamoxifen was administered to adult mice (aged 8-20 weeks) via intraperitoneal injections. For each injection, mice received 50 mg/kg body weight of TX (Sigma-Aldrich) dissolved in 100% corn oil (Sigma). Mice were usually given daily TX injections for five consecutive days, but in some experiments, this was reduced to injections for 3 consecutive days. “No TX” controls were given only vehicle (corn oil). All animal experiments were conducted under a protocol approved by the Institutional Animal Care and Use Committee (IACUC) of the University of South Carolina. Both male and female mice were included in the experiments, with no observed differences in outcomes between sexes.

### Tissue Collection and Immunohistochemistry

Animals under deep isoflurane anesthesia were transcardially perfused with ice-cold PBS, followed by 4% paraformaldehyde (PFA) in 0.1 M PBS (pH 7.4). Tissue samples from the brain, spinal cord, and sciatic nerve were harvested and fixed in 4% PFA in PBS overnight. Tissues were then washed in phosphate buffered saline (PBS), cryoprotected in 30% w/v sucrose in PBS, embedded in Optimal Cutting Temperature Compound (O.C.T., Fisher) and stored at −80°C. Tissues were cryo-sectioned at thicknesses ranging from 10 to 50 μm, and stored at −80°C. For immunostaining, sections were rehydrated in PBS, permeabilized with 0.3% Triton X-100 for 10–30 minutes, blocked with 10% goat serum, 3% BSA, and 0.1% Triton X-100 in PBS for 1 hour, then incubated with primary antibody overnight at 4°C. Primary antibodies used are as follows: LIS1 mouse monoclonal (Santa Cruz Biotechnology Cat# sc-17817, RRID:AB_627889; IF: 1:200, WB: 1:1000). NFL, NFM, and NFH, neurofilament light, medium, and heavy chain chicken polyclonals. Aves, RRIDs: AB_2313553, AB_2313554, AB_2313552; IF: 1:500); NF 200-kDa mouse monoclonal, clone RT97 (Millipore CBL212, RRID:AB_93408; IF: 1:500); EEA1 rabbit monoclonal antibody (Cell signaling Technology C45B10, RRID:AB_2096811); AIF XP rabbit monoclonal antibody (Cell signaling Technology #5381, RRID:AB_10634755); Degenotag mouse monoclonal antibody to NF light chain (EnCor Biotechnology, Cat# MCA-1D44, RRID:AB_2923483; IF 1:1000); dynein intermediate chain mouse monoclonal (Santa Cruz Biotechnology sc-13524, RRID:AB_668849; IF: 1:200 WB: 1:1000) Vglut1 mouse monoclonal antibody (Invitrogen Cat # PIPA585764; IF: 1:250). GFAP mouse monoclonal antibody (Santa Cruz Biotechnology Cat # sc-33673, RRID:AB_627673; IF 1:1000, WB 1:500); After three PBS washes, sections were incubated for 2 hours with secondary antibodies (Alexa Fluor 555-or 647-conjugated goat anti-mouse (Invitrogen, A21422, 11001); Cy5-conjugated donkey anti-chicken, mouse, goat (Jackson ImmunoResearch 703-175-155, 715-175-150, and 705-175-147, RRIDs: AB 2340365, AB 2340819, AB 2340415. Nuclei were stained with Hoechst dye or DAPI, and samples were mounted using Prolong Gold. Cryosections were imaged using a Leica STELLARIS 5 confocal microscope with LAS X software and a 20× or 63× oil immersion objective (1.4 N/A) or a Zeiss Axiovert 200 inverted microscope equipped with AxioVision software, a Plan-Neofluor 20× dry objective and Plan-Neo 100×/1.30 or Plan-Apo 63×/1.40 oil-immersion objectives (Immersol 518F; Carl Zeiss, Inc.)

### Counting of axonal swellings and fragmentation in tissue sections, teased nerves and cultured neurons

Axonal swellings were quantified in tissue sections and axons in DRG cultures. A swelling was counted if it had a diameter at least twice that of the associated axon. Swellings are labeled intensely with EYFP, probably due to accumulation. Fragmentation was determined for teased nerve preparations if the EYFP/tdTomato signal was broken along the length of the axons.

### Fluoromyelin Staining

Tissues were rehydrated with PBS 3 times for 5 minutes each. Fluoromyelin red (Invitrogen, Cat# F34652) was diluted to 1X and applied for 20 minutes at room temperature. Tissues were rinsed 3 times for 5 minutes with PBS followed by one rinse with DI water. Tissues were air-dried at room temperature for 5 minutes and mounted with ProLong Gold antifade reagent with DAPI (Invitrogen, Cat# P36935). Tissues were imaged using a Zeiss AxioObserver 7 with an Apotome 3. Images were acquired with a Plan-Apochromat 20x/0.8 M27 air objective. 5 apotome images and optical sections at 0.2 um z step size were taken per field of view.

### Protein isolation and immunoblotting

Tissues were rapidly dissected from CO2-euthanized mice and immediately frozen in liquid nitrogen, followed by homogenization using a Dounce homogenizer in ice-cold RIPA lysis buffer (50 mM Tris-HCl, pH 8.0, with 150 mM sodium chloride, 1.0% NP-40, 0.5% sodium deoxycholate, and 0.1% sodium dodecyl sulfate) containing HALT protease and phosphatase inhibitors (ThermoFisher, Cat# 78440). Protein concentrations in the extracts were quantified using a BCA assay (ThermoFisher, Cat# 33227). For Western blotting, 5 µg of Brain and spinal cord samples were separated on 4-15% acrylamide gels (BioRad, Cat # 4561084) and transferred to PVDF membranes. The blots were probed with antibodies against LIS1 and dynein intermediate chain, with protein detection carried out via chemiluminescence using Secondary antibodies used: HRP-conjugated goat anti-rabbit and mouse (Millipore 12-348 and 12-349, RRIDs:AB_390191 and AB_390192). All blots are representative of at least three independent experiments.

### Teased nerve preparation

Animals were deeply anesthetized with isoflurane and perfused transcardially with ice-cold PBS, followed by 4% PFA in 0.1 M PBS. The sciatic nerve was exposed by separating the biceps femoris along the intermuscular septum, then cut at the knee and sciatic notch before being fixed in 4% PFA overnight at 4°C. After post-fixation, a 3-5 mm nerve segment was trimmed and placed in PBS. Under a stereomicroscope, the epineurium was removed, and the nerve was teased into individual fibers on an adhesive slide. The slide was air-dried and stored at 4°C overnight before immunostaining the fibers for analysis.

### Preparation of astrocyte cultures

Cultures were generated from iKO mice (TX-*Aldh1l1-LIS11 ^fl/fl^)* and control mice TX*-Aldh1l1-LIS11 ^wt/wt^* or TX*-LIS1^fl/fl)^*. Mice were euthanized on day 10 after the initial injection, then cortices removed and placed in cold Hanks Buffered Saline Solution (HBSS). After removing meninges and cutting cortices into small pieces, pieces were washed in HBSS 3 times and subjected to trypsin dissociation at 37°C for 20 minutes. Cells were dissociated by trituration with a flamed Pasteur pipet, then incubated at 37°C for 10 minutes. Cells were triturated again, then passed through a70 µm filter. Cells were pelleted by centrifugation at 8000 x g for 10 minutes. Pellets were resuspended in 10 mL of astrocyte media (DMEM/F12, 5% fetal bovine serum, 0.5% Non-essential Amino Acids and 0.5% penicillin-streptomycin), plated into T25 flasks, and maintained at 37°C, 5% CO2. Medium was refreshed every 48 hours for 8 days, then cells placed on a shaker to shake at 37°C for 2-4 hours to remove microglia. For imaging cells to were plated onto poly-D-lysine coated 12mm coverslips. Cells were fixed with 4% paraformaldehyde, processed for immunofluorescence, then mounted onto slides for confocal or widefield microscopy.

### Preparation of DRG cultures

Mice were injected with TX in three daily doses. Dissociated cultures of adult DRG from iKO mice (TX-*Thy1-LIS1^fl/fl^*) and control mice CreERT2 (TX-*LIS1^fl/fl^* or TX-*Thy1-LIS1^wt/wt^*) were prepared as described by Twiss et al. (2000). DRGs were harvested in Hybernate A medium (ThermoFisher, Cat# A1247501) and dissociated using Collagenase type 2 (ThermoFisher, Cat#17101015) at 37°C with 5% CO2 for 15 minutes. DRGs were then triturated using a fire-polished Pasteur pipette, diluted in 5 ml DMEM/F12 (ThermoFisher, Cat#MT16405CV), and centrifuged at 100 x g for 5 minutes. After pelleting, dissociated DRGs were washed in DMEM/F12 and cultured in DMEM/F12 supplemented with 1x N1 supplement (Sigma, Cat#N6530), 10% fetal bovine serum (Hyclone, Cat#SH3007102), and 10 µM cytosine arabinoside (Sigma, Cat# C6645). The cells were plated on poly-L-lysine (ThermoFisher, Cat#A005C) and laminin (Sigma, Cat#CC095)-coated glass coverslips. After 72 hours, DRG cultures were harvested for immunostaining and imaging.

### Experimental design and statistical analysis

Statistical tests and data graphing were performed using GraphPad Prism version 10. Individual animals were considered as biological replicates (*n*), and all experiments involve comparisons to littermate controls. The number of animals used (*n*), statistical measures represented in graphs, statistical tests performed, and *p* values are all reported in the legends. The threshold for statistical significance was defined as *p* < 0.05.

## RESULTS

### LIS1 depletion exclusively in astrocytes does not explain the phenotypes observed in TX*-Act-LIS1^fl/fl^* mice

Astrocytes have important regulatory roles in the nervous system. They provide metabolic support to neurons, modulate synaptic function and maintain the blood brain barrier. Astrocytes also contribute to circuit remodeling by synapse phagocytosis (Chung et al., 2024), a process that involves both LIS1 and dynein (Chhatre et al., 2020; Omer et al., 2024). If LIS1 depletion disrupts one or more of these processes in adult astrocytes it could potentially lead to neurological impairment.

To ensure that LIS1 is expressed in adult astrocytes we examined dissociated cultured astrocytes using LIS1 immunofluorescence (IF). This allowed better separation of astrocytes from other cells in the brain. Astrocytes were harvested from adult mouse cortices and plated onto poly-D-lysine and laminin coated coverslips for staining and visualization. LIS1 displayed a typical punctate distribution with enrichment at centrosomes **(Fig. 1 A)**. Dynein was also present in cultured astrocytes and enriched at centrosomes **(Fig. 1B**). LIS1 was also frequently enriched around what appear to be pyknotic nuclei that were likely phagocytosed from cells that died during the isolation procedure **(Fig. 1 C, D).** This confirms the presence of LIS1 in adult mouse astrocytes, where it may contribute to the phagocytosis of cellular debris (Chhatre et al., 2020; Lee et al., 2021).

**Figure 1:**
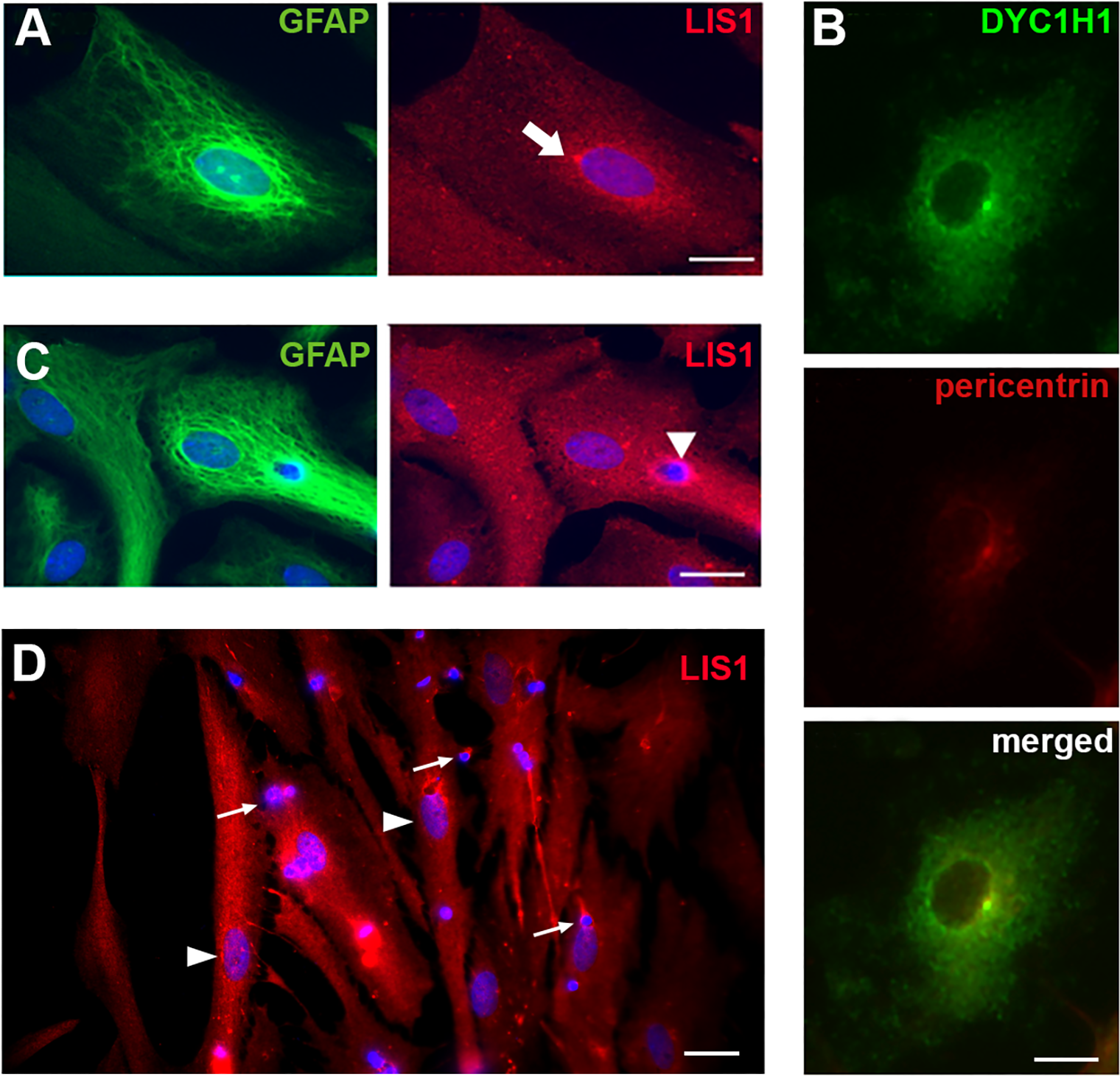
LIS1 and dynein distribution in cultured adult mouse astrocytes. **A** LIS1 immunofluorescence (red) is prominent at the centrosome (arrow) near the nucleus stained with DAPI (blue) in an interphase astrocyte also positive for GFAP (green). **B)** Dynein labeled with a dynein intermediate chain antibody (green) is also enriched at a centrosome, here labeled with the centrosomal marker, pericentrin (red). **C)** LIS1 is enriched around an engulfed pynknotic nucleus (arrowhead). **D)** Pynknotic nuclei likely arise from neurons or other cell types that do not survive the dissociation process (arrows). In this image two healthy astrocyte nuclei are indicated with arrowheads. Scale bars = 10 µm.

To determine the importance of LIS1 expression in adult mouse astrocytes we used an *Aldh1l1* promoter to drive expression of CreERT2 specifically in astrocytes, generatingTX*-Aldh1l1-LIS1^fl/fl^* mice. These were crossed with a reporter strain that expresses tdTomato in cells with activated CreERT2 recombinase **(Fig. 2A)**. We used a standard IP injection TX dosing regimen, administering 50 mg/kg TX once a day for 5 days to both TX*-Aldh1l1-LIS1^fl/fl^* mice and TX*-Aldh1l1-LIS1^wt/wt^* controls **(Fig. 2A)**. Despite strong induction of tdTomato in all astrocytes by day 10 after the initial injection **(Fig. 2B)**, TX-*Aldh1l1-LIS1^fl/fl^* mice show none of the neurological impairments seen in the TX-*Act-LIS1^fl/fl^* mice (Hines et al., 2018). Astrocytes cultured from adult TX-*Aldh1l1-LIS1^fl/fl^* mice had reduced LIS1 expression **(Fig. 2C).**

**Figure 2:**
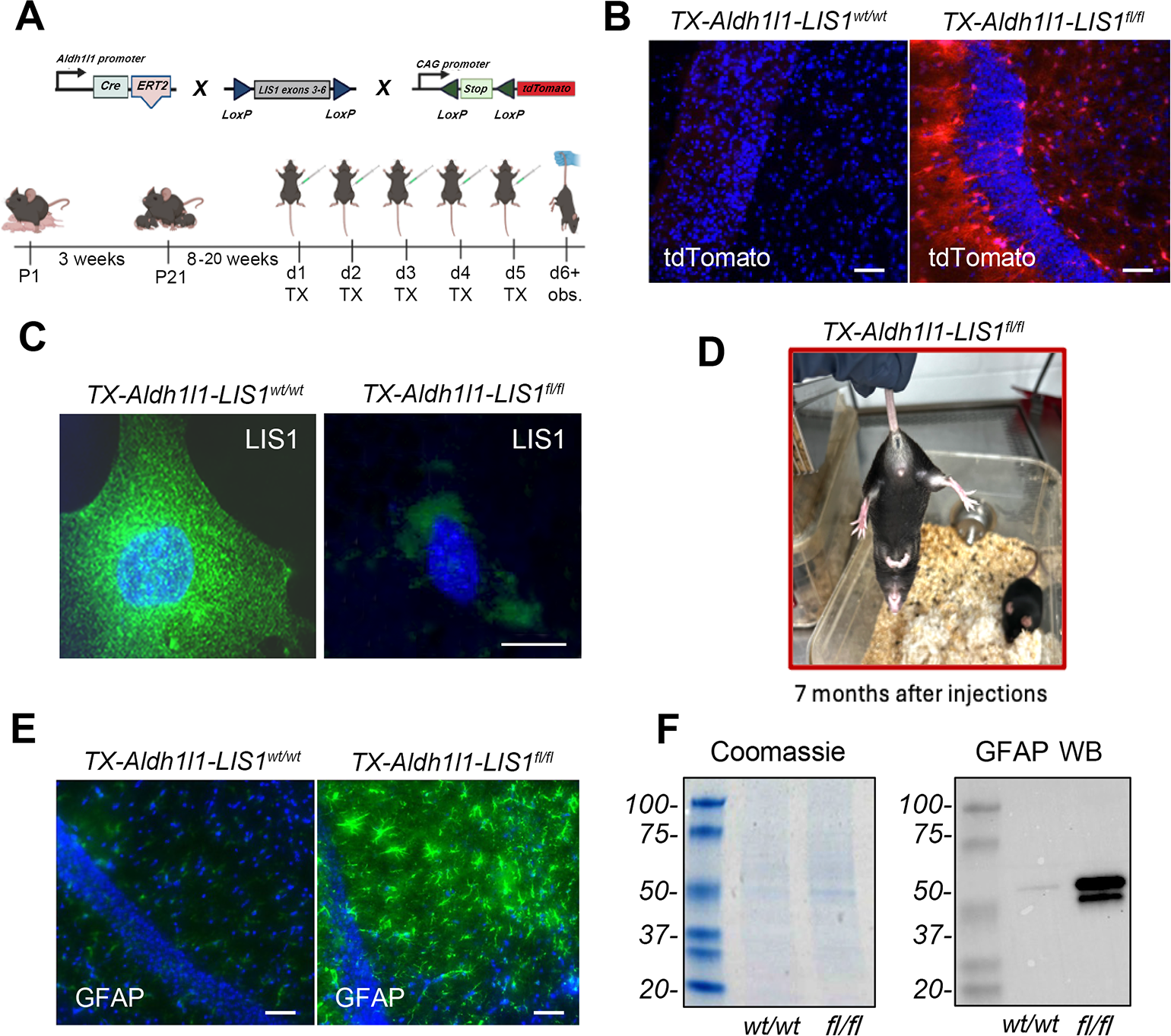
TX-*Aldh1l1-LIS1^fl/fl^* mice do not phenocopy TX-*Act-LIS1^fl/fl^* mice, but astrocytes show elevated GFAP levels. **A)** Top diagram: Schematic of mouse crosses used to generate astrocyte specific iKO of LIS1 (*Aldh1l1-LIS1^fl/fl^)*. CreERT2 is a fusion of CreERT2 recombinase and a modified ligand-binding domain of the estrogen receptor, which is activated by the synthetic ligand tamoxifen (TX). CreERT2 mice are crossed with floxed LIS1 mice, where exons 3-6 of the LIS1 gene are flanked by LoxP sites and excised upon CreERT2 activation. Bottom diagram: TX is administered to adult mice (8-20 weeks old) at 50 mg/kg via IP injection for 5 consecutive days. **B)** tdTomato CreERT2 activity reporter is increased in astrocytes of in *TX- Aldh1l1-LIS1^fl/fl^* mice. Scale bar = 25 µm. Blue = DAPI in all images. **C)** LIS1 expression (green) is reduced in astrocytes cultured from adult *TX- Aldh1l1-LIS1^fl/fl^*mice compared to *TX- Aldh1l1-LIS1^wt/wt^* control mice. Scale bar = 10µm. N≥3 culture preparations. **D)** *TX- Aldh1l1-LIS1^fl/fl^* mice do not develop neurological impairments and survive at least 7 months after the first TX injection. **E)** Representative brain section stained with GFAP (green) and DAPI (blue), showing upregulation of GFAP expression in astrocytes of *TX- Aldh1l1-LIS1^fl/fl^* mice. Scale bar = 10 µm. N≥ 4 mice per group. **F)** Western blot of brain tissue confirms increased GFAP expression in *TX- Aldh1l1-LIS1^fl/fl^* mice. A Coomassie-stained gel shows equal protein loading.

Dynein moves a variety of cellular components towards the centrosome in cells with a radial MT array, and disruption of dynein leads to organelle dispersion (Reck-Peterson et al., 2018). We reported that organelles and dynactin appeared more diffuse and less concentrated at centrosomes in mouse embryo fibroblasts (MEFs) harvested from LIS1^+/-^ embryos, an early indication that LIS1 loss might negatively impact dynein processivity (Smith et al., 2000).To visualize how dynein is impacted by LIS1 depletion in adult mouse astrocytes we examined its distribution using an antibody against dynein intermediate chains. As with the MEFs, more diffuse dynein was observed in astrocytes with reduced LIS1 expression harvested from TX-*Aldh1l1-LIS1^fl/fl^* mice **(Fig. S1A).**

Increased dynein/dynactin/cargo accumulation at centrosomes has been used as a readout for increased dynein processivity (Schroer et al., 1989; King and Schroer, 2000; Kapitein et al., 2010; Gao et al., 2015; Wang et al., 2019; Christensen et al., 2021). LIS1 overexpression causes these changes, and we interpret this as evidence for dynein stimulation due to increased LIS1 protein relative to dynein (Smith et al., 2000; Gao et al., 2015). Given that LIS1 binding prevents dynein from transitioning into an autoinhibited phi-like conformation, we suspect that increasing the ratio of LIS1 to dynein increases the proportion of non-phi, activatable motors. The change in steady-state distribution of dynein to centrosomes makes sense given the high concentration of MT minus ends at this location. We therefore measured the fluorescence intensity of dynein in 30 TX-*Aldh1l1-LIS1*^wt/wt^ and 30 TX-*Aldh1l1-LIS1^fl/fl^*astrocytes from 3 different animals in ImageJ, using a circle ROI to measure average pixel intensities at centrosomes labeled with pericentrin or **γ**-tubulin. There is a significant reduction in dynein intensities in LIS1 depleted astrocytes pointing to possible reduced numbers of processive motors **(Fig. S1B).**

Dynein contributes to MT organization in cells via MT sliding (Shima et al., 2006; Tanenbaum et al., 2013; Bender et al., 2015; Rao et al., 2017). In neurons, dynein drives MTs out into axons (Ahmad et al., 1998; He et al., 2005). We reported that LIS1 overexpression in Cos-7 cells caused MTs to move to the cell periphery and found that MTs were more tightly clustered around the nucleus in LIS1^+/-^ MEFS (Smith et al., 2000). To determine if MT distribution was altered similarly in LIS1-depleted astrocytes we stained cells using an **α**-tubulin antibody and utilized a segmented line tool in ImageJ to draw a line of equal width (2.5 µm) around the outside of nuclei labeled with DAPI. Average pixel intensity was determined for 30 cells for each treatment. As with the LIS1^+/-^ MEFs, tubulin IF intensity was significantly increased in LIS1-depleted cells (**Fig. S1 C, D).** Together these experiments demonstrate that LIS1 expression is reduced in *Aldh1l1-LIS1^fl/fl^* astrocytes, and that this impacts normal dynein and MT distribution. Despite these cellular changes, TX-*Aldh1l1-LIS1^fl/fl^* mice live normal life spans with no obvious neurological deficits such as leg clasping upon tail suspension **(Fig. 2D)**. However, astrocytes throughout the brain and spinal cord of TX-*Aldh1l1-LIS1^fl/fl^* mice displayed a strong increase in GFAP expression **(Fig. 2E, F)**. This was also observed in the TX-*Act-LIS1^fl/fl^* mouse model(Hines et al., 2018). Our new finding indicates that this is a cell autonomous response to LIS1 loss in astrocytes rather than a response triggered by alterations in neighboring LIS1-depleted neurons. Future studies will determine if loss of LIS1 in astrocytes produces more subtle behavioral consequences in mice due to the cellular changes in GFAP, dynein and MTs.

### LIS1 depletion in projection neurons results in the rapid onset of neurological impairments and death

To determine the importance of LIS1 expression specifically in projection neurons we used the SLICK-H mouse strain to express CreERT2(Vidal et al., 1990; Feng et al., 2000; Young et al., 2008). Two separate Thy1 promoters controlled the expression of both an EYFP protein to identify projection neurons and the Cre recombinase enzyme **(Fig. 3A).** We used the same dosing regimen for these mice as with the TX-*Aldh1l1-LIS1* mice, 50 mg/kg TX once a day for 5 days **(Fig. 3B)**. All projection neurons in SLICK-H mice are all labeled with EYFP, regardless of whether CreERT2 is activated by TX **(Fig. 3C** shows cortical pyramidal neurons). N either TX-*Thy1-LIS1^wt/wt^* control mice nor heterozygous iKO mice (TX-*Thy1-LIS1 ^fl/wt^*) show any overt neurological phenotypes **(Movie 1)**. Homozygous iKO mice (TX-*Thy1-LIS1^fl/fl^*), have reduced LIS1 expression compared to controls, both by western blotting and IF **(Fig. 3D, E).** These mice displayed leg clasping upon tail suspension on day 6 after the first injection **(Fig. 3B, F)**. More severe symptoms became apparent between day 9 and 10 after the first injection **(Movie 2, Fig. 3B)**. Mice had an altered gait, exhibited severe shivering and “Straub tail” a reaction caused by contraction of the sacrococcygenus muscle (Bilbey et al., 1960). There was also significant weight loss in the iKO mice. Homozygous iKO mice were routinely euthanized on day 10 after the initial TX injection.

**Figure 3:**
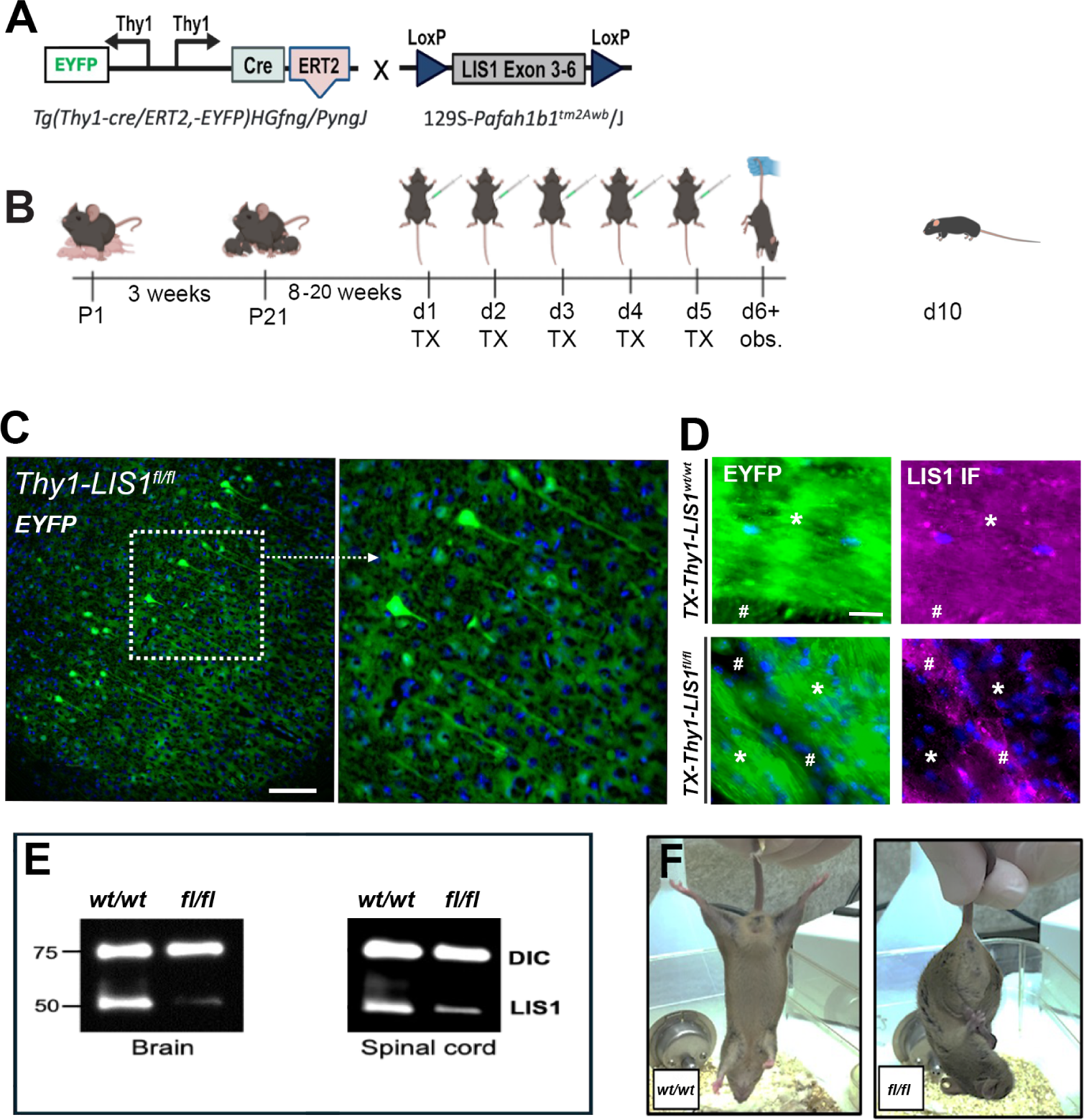
TX-*Thy1-LIS1^fl/fl^*mice have reduced LIS1 in fiber tracts and develop severe neurological symptoms. **A)** Schematic illustration of the generation of projection neuron specific conditional LIS1 knockout mice (TX-*Thy1-LIS1^fl/fl^*). Two copies of the Thy1 promoter separately drive the expression of EYFP and the inducible CreERT2. **B)** The timeline of TX administration is similar to that of the TX-Aldh1l1-LIS1 mice, but iKO in projection neurons caused severe neurological symptoms and mice were euthanized on day 10 after the first injection. **C)** All projection neurons are labeled with EYFP (green) regardless of CreERT2 recombinase activity. Shown are labeled neurons in the cerebral cortex. Scale bar = 10 µm. The zoomed shows clear EYFP signal in axons and cell bodies. Blue = DAPI in all images. EYFP-labeled axon tracts running in brainstem sections were processed for LIS1 immunofluorescence (magenta). The regions marked with an asterisk show that LIS1 expression is reduced in EYFP-labeled fiber tracts in *TX-Thy1-LIS1^fl/fl^* mice compared to control *TX-Thy1-LIS^wt/wt^* mice. The hashtags show that regions not positive for EYFP have substantial LIS1 labeling in both *TX-Thy1-LIS1^fl/fl^*and *TX- Thy1-LIS^wt/wt^* mice. Scale bar = 50 µm. N≥ 4 mice per group. **D)** Reduced LIS1 expression in *TX-Thy1-LIS1^fl/fl^* mice (*fl/fl*) compared *TX-Thy1-LIS^wt/wt^* mice (*wt/wt*) confirmed by western blotting of extracts from brain and spinal cord prepared on day 10. Levels of dynein intermediate chain (DIC) are not reduced and serve as a loading control. **E)** On day 6 after the initial injection *TX-Thy1-LIS1^fl/fl^* mice exhibit neurological phenotypes such as leg clasping upon tail suspension, shown here.

### LIS1 depletion in projection neurons results in axonal pathologies

Axons in *TX-Thy1-LIS1^fl/fl^* mice but not *TX-Thy1-LIS1^wt/wt^* controls developed axonal swellings by day 10 after the initial injection, as shown here in longitudinal spinal cord sections (**Fig. 4A, B**). The number of swellings was significantly higher in *TX-Thy1-LIS1^fl/fl^* mice compared to controls **(Fig. 4C)**. Additionally, the number of swellings in the spinal cord was progressive, being significantly higher on day 6 after the initial injection than on day 5, which correlates with the onset of neurological symptoms on day 6 **(Fig. 4D)**. The swellings and fragmentation are reminiscent of the degeneration of axons distal to the site of a nerve injury (Wallerian degeneration). Similar axonal pathologies are also typical of most if not all neurodegenerative diseases, and indeed, may involve similar pathways that trigger programmed axonal death(Coleman and Hoke, 2020).

**Figure 4:**
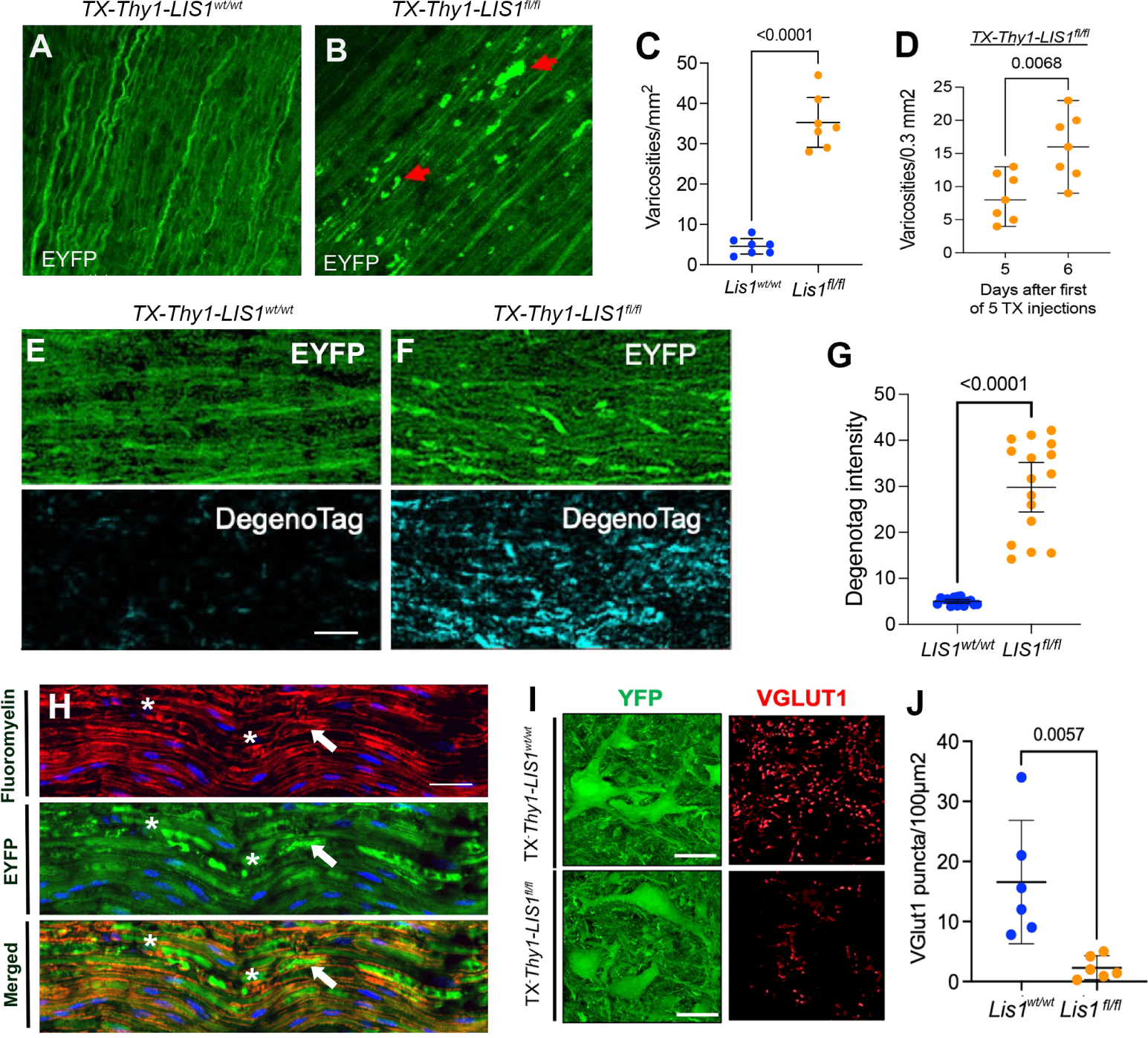
LIS1 depletion in projection neurons causes axonal degeneration and synapse loss. **A, B)** Representative images of EYFP-positive axons in frozen longitudinal spinal cord sections from *TX- Thy1-LIS^wt/wt^* control mice *(LIS1^wt/wt^*; left) and *TX-Thy1-LIS1^fl/fl^* iKO mice (*LIS1^fl/fl^*; right) mice treated with 5 doses of TX and perfused on day 10 after the initial injection. Axonal swellings (red arrows) are apparent in the *LIS1^fl/fl^*spinal cord. Scale bar = 50 µm. n ≥ 6 mice per group. **C)** Significantly more varicosities per mm² are present on day 10 in LIS1 depleted spinal cords compared to control spinal cords. Graph shows the mean ± 95% CI. Significance determined by two-tailed Student’s t test for 7 biological replicates. **D)** After 5 daily doses of TX, there were more varicosities *LIS1^fl/fl^* spinal cord on day 6 than on day 5, demonstrating early progression. N ≥ 3 mice per group. **E-G)** Representative images of EYFP and Degenotag IF in longitudinal spinal cord sections from control *TX-Thy1-LIS^wt/wt^* (left) and *TX- Thy1-LIS1^fl/fl^*(right) mice perfused on day 10. Scale bar = 50 µm. Graph shows the mean ± 95% C. Significance determined by two-tailed Student’s t test two biological replicates, 8 regions per tissue section. **H)** Myelin labeled with fluoromyelin in TX-*Thy1-LIS1^fl/fl^* sciatic nerve section appears fragmented and has accumulated in some regions (asterisks) but does not always correlate with sites of EYFP accumulation in axons (arrow). Scale bar = 25µm. **(I, J)** Synapses in the spinal cord ventral horn labeled with VGLUT1 (red) in TX-*Thy1-LIS1^fl/fl^* mice. A reduction in VGLUT1-labeled synapses was observed, suggesting that LIS1 depletion may disrupt synaptic integrity, potentially secondary to axonal degeneration. Quantification shows the number of VGLUT1 puncta per 100 µm. Graph shows the mean ± 95% C for 3 biological replicates 3 sections per mouse. Scale bar = 40 µm. Significance determined by two-tailed Student’s t test.

To definitively assess whether the fragmentation represents axon degeneration, we used the “Degenotag” antibody for immunofluorescence on spinal cord sections. This antibody has been used in the study of degenerative diseases and after nerve injury, as it allows visualization of neurofilament (NF) proteolysis by calpain and other proteases (Shaw et al., 2023). The Degenotag antibody recognizes a neurofilament light chain epitope not exposed until the protein is proteolytically cleaved. Spinal cord sections from TX-*Thy1-LIS1^fl/fl^*mice exhibited substantially higher Degenotag immunofluorescence than those from TX-*Thy1-LIS1^wt/wt^* controls, further confirming active axonal degeneration in the LIS1 depleted neurons **(Fig. 4E-G).** In Wallerian degeneration, the myelin sheath also breaks down in the nerve distal to the injury site (Conforti et al., 2014). To determine if there is myelin breakdown in sciatic nerve sections from TX-*Thy1-LIS1^fl/fl^*mice we used fluoromyelin staining **(Fig. 4H)**. There appears to be fragmentation and accumulation of myelin, supporting the idea that LIS1 depletion in projection neurons induces Wallerian-like degeneration. Future studies will be needed to determine if there is evidence of Schwann cell changes or macrophage invasion in nerves undergoing degeneration due to LIS1 depletion, as this is typical of Wallerian degeneration(Zhao et al., 2022). Because axons were degenerating in TX-*Thy1-LIS1^fl/fl^* mice, we reasoned that synapses may also be impacted. Indeed, VGLUT1 immunofluorescence revealed fewer synapses on the somatic motor neurons in the spinal cord ventral horns of TX-*Thy1-LIS1^fl/fl^* mice compared to controls on day 10 after initial TX exposure **(Fig. 4I, J).** We have not detected apoptosis of LIS1-depleted neurons on day 10 after 5 injections, indicating that LIS1 depletion may trigger axonal death without causing neurons themselves to die.

### Widespread neuronal Cre activation correlates with axon fragmentation in sciatic nerves of TX-*Thy1-LIS1^fl/fl^* mice

*TX-Thy1-LIS1^fl/fl^* mice were crossed with tdTomato Cre activity reporter mice **(Fig. 5A).** Virtually all EYFP-positive neurons in the hippocampus of *TX-Thy1-LIS1^fl/fl^*mice also had tdTomato fluorescence by day 6 after the initial TX injection **(Fig. 5B, C)** and this was true throughout the brain and spinal cord (not shown) as well as in peripheral nerves **(Fig. 5D).** Control mice without Thy1-CreERT2 expression did not express tdTomato, other than an occasional cell, as indicated in the original paper describing these mice (Young et al., 2008). Axons labeled with both EYFP and tdTomato in teased sciatic nerve preparations from *TX-Thy1-LIS1^fl/fl^* mice, but not *TX-Thy1-LIS1^wt/wt^*controls, were often fragmented and showed swellings like those observed in spinal cord tissue sections (**Fig. 5D, E).** Both sensory and motor axons developed swelling and fragmentation, as detected in teased nerve preparations of sural (sensory) and peroneal (motor) nerves **(Fig. 5F)**.

**Figure 5:**
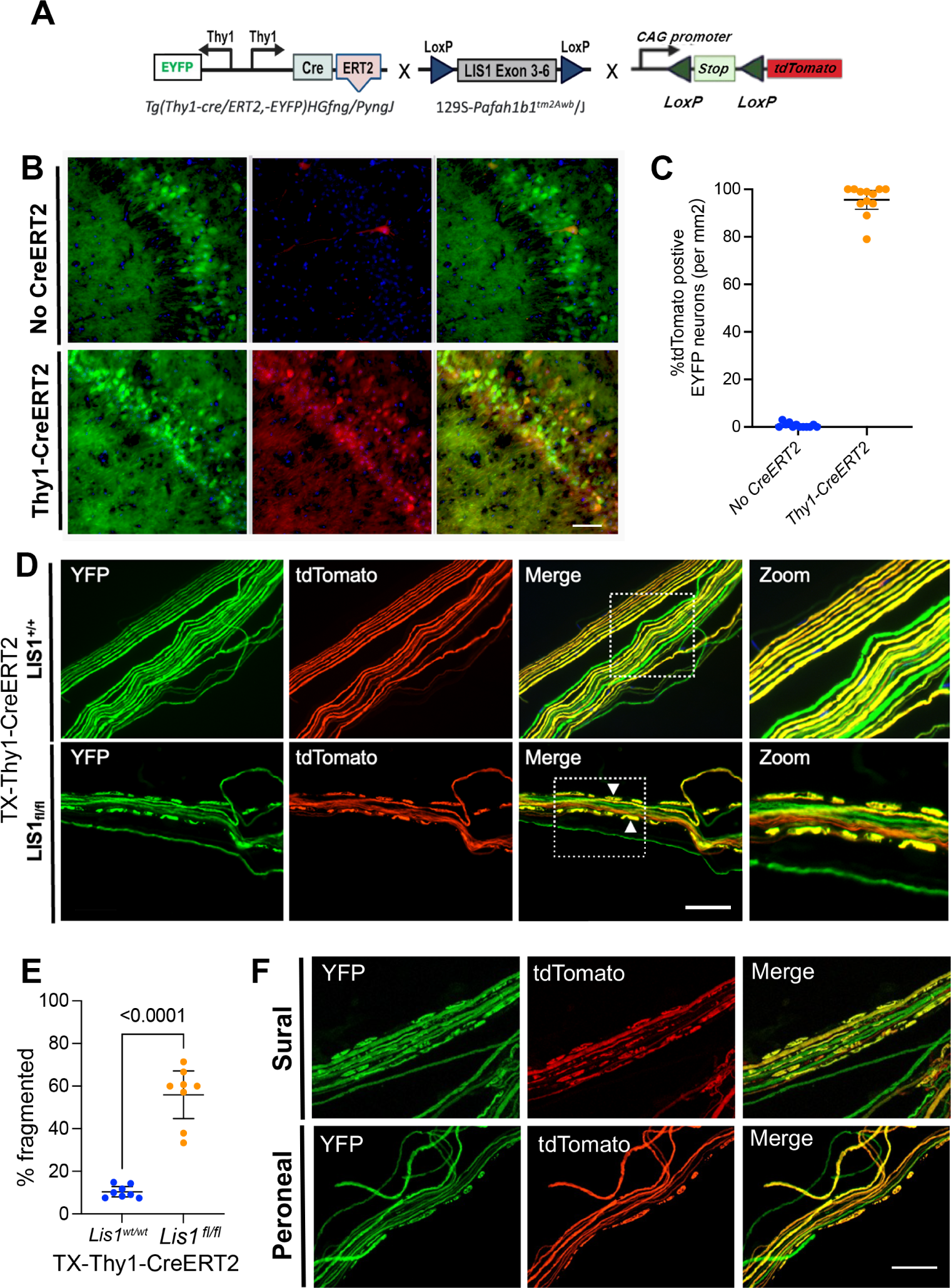
Axonal degenerative profiles correlate with CreERT2 activity. **A)** Thy1-CreERT2 mice (both with and without floxed LIS1 alleles) were crossed with the tdTomato Cre activity reporter strain. **B)** tdTomato-positive neurons overlap substantially with EYFP positive neurons in the hippocampus on day 10 after the initial TX injection. Scale bars = 10 µm. **C)** The percent overlap was statistically significant compared to no-CreERT2 controls, which had a very small number of tdTomato positive cells even in the absence of TX. Graph shows the mean ± 95% C for 3 biological replicates, 3 sections per mouse. Significance determined by two-tailed Student’s t test. **D, E)** Individual axons in teased sciatic nerves are readily distinguishable and labeled with both EYFP and tdTomato. Only axons from TX-*Thy1-LIS1^fl/fl^* nerves showed significant levels of fragmentation. Graph shows the mean ± 95% C for 2 biological replicates, 4 regions of the teased nerve per sample. Significance determined by two-tailed Student’s t test. **F)** Axons in teased nerve preparations from the sural (sensory) and peroneal (motor) nerves of *TX-Thy1-LIS1^fl/fl^* mice show axonal fragmentation. Scale bars = 50 µm; Scale bar in zoomed images in D = 100µm.

### *TX-Thy1-LIS1^fl/fl^* exhibit fewer neurological phenotypes after only 3 TX injections

We tested 12 mice *TX-Thy1-LIS1^fl/fl^* and 12 *TX-Thy1-LIS1^wt/wt^* controls with a lower TX dose by giving 3 consecutive daily TX injections instead of 5. After only 3 TX injections LIS1 iKO mice survived beyond day 10 and showed no shivering, seizures, or weight loss at that time **(Fig. 6A, B)**. Upon examination of brain sections, it became clear that fewer neurons had activated CreERT2 in mice that only received 3 TX injections which likely explains the less severe phenotypes **(Fig. 6C, D)**. Mice did exhibit leg clasping upon tail suspension on day 6 and began to have increased neurological symptoms. Mice were euthanized 22 days after the initial injections. Because animals survive longer with 3 injections, we were able compare axonal swelling on day 10 and day 22 in these mice. Interestingly there are significantly more swellings on day 22 after the initial injection **(Fig 6. E, F**). These findings indicate that there is a correlation between symptom severity and number of neurons with LIS1 loss. They also add support to the idea that LIS1 depletion initiates a progressive axonal degeneration process that continues to worsen over time, in that neurons with LIS1 depletion will continuously degenerate if the animal’s survival is sustained by the unaffected neurons.

**Figure 6:**
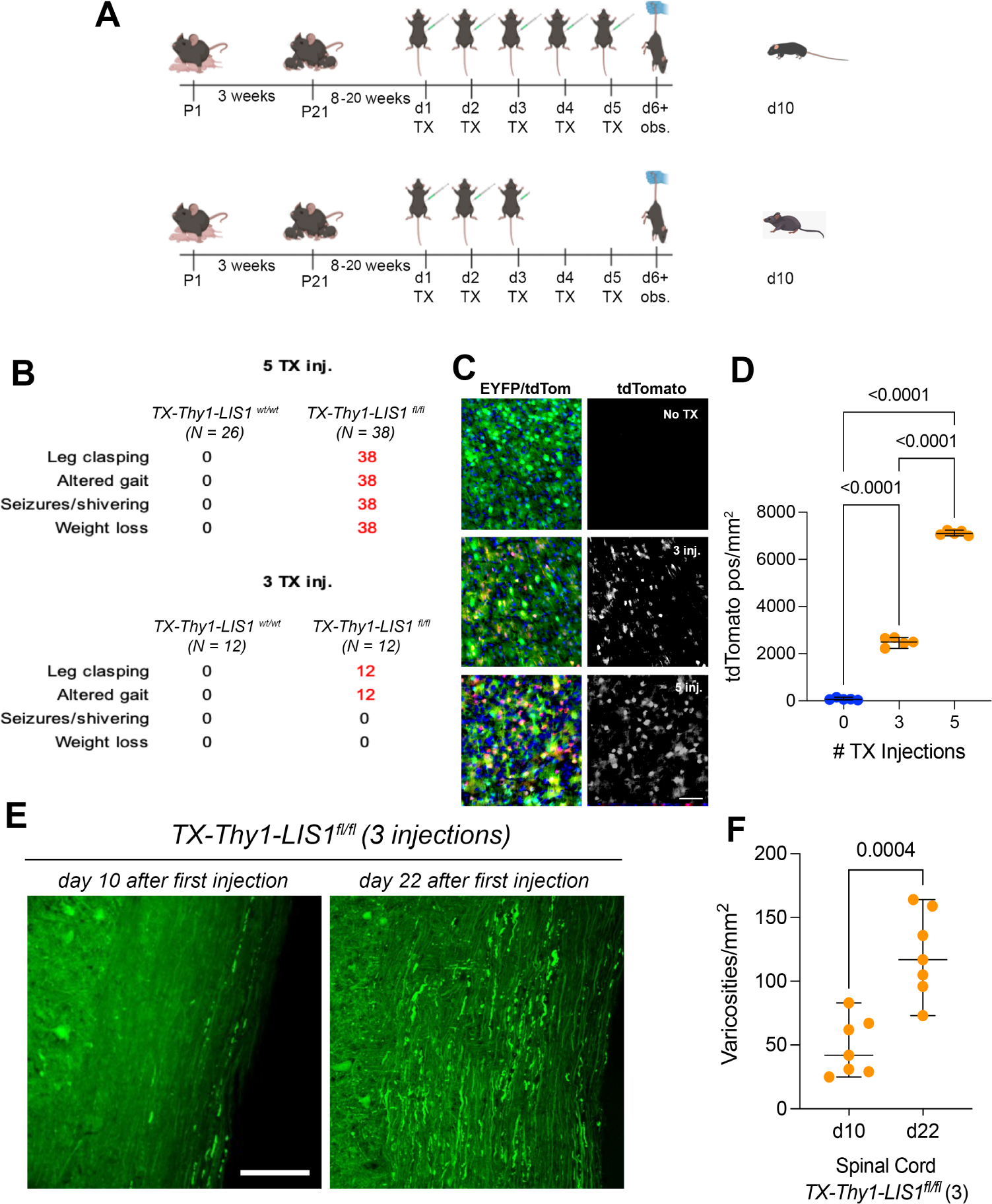
Fewer TX injections activate CreERT2 recombinase in fewer neurons resulting in fewer symptoms and delayed death. **A)** We compared the effects of giving 3 or 5 daily TX injections to TX-*Thy1-LIS1^fl/fl^* and TX*-Thy1-LIS1^wt/wt^* mice. Phenotypes were assay on days 6, 10 and 22 after the first injection. All TX-*Thy1-LIS1^fl/f^* that received 5 injections were euthanized on day 10 after the first injection. **B)** The number of mice that displayed the indicated symptom is shown for each genotype. Seizures and weight loss were only observed in TX-*Thy1-LIS1^fl/f^* mice that received 5 injections*. TX-Thy1-LIS1^fl/^*^f^ mice that received only 3 injections had the leg clasping phenotype, but survived for 22 days, at which time they were euthanized. **C)** After only 3 injections fewer neurons in the cortex on day 10 after the initial injection were positive for the tdTomato CreERT2 activity reporter. Scale bar = 100 µm. **D)** Graph shows mean ± 95% CI for five biological replicates. Significance determined using ordinary one-way ANOVA with multiple comparisons. **E, F**) Axons in TX-*Thy1-LIS1^fl/fl^* mice that received 3 days of TX also showed degenerative swellings, and the number increased between day 10 and 22 after the first injection. Scale bar = 50 µm. **F)** Quantitation of varicosities per mm² in the spinal cord on day 10 or day 22 after the first injection Graph shows mean ± 95% CI for three biological replicates ≥ 2 sections per animal. Significance determined using a two-tailed Student’s t test.

### LIS1 depletion does not prevent axon growth in culture, but axons show signs of degeneration and cargo distribution changes

Cultured DRG neurons from *TX-Thy1-LIS1^fl/fl^* animals exposed to 3 TX injections were able to extend long axons in culture despite showing reduced LIS1 staining, particularly in growth cones and axon shafts **(Fig. 7A).** They were also positive for Degenotag, indicating that despite axon extension, pathological changes to NF light chains were present. **(Fig. 7B)**. In addition, these axons appeared more fragile than controls and exhibited increased numbers of large varicosities **(Fig. 7C, D).**

**Figure 7:**
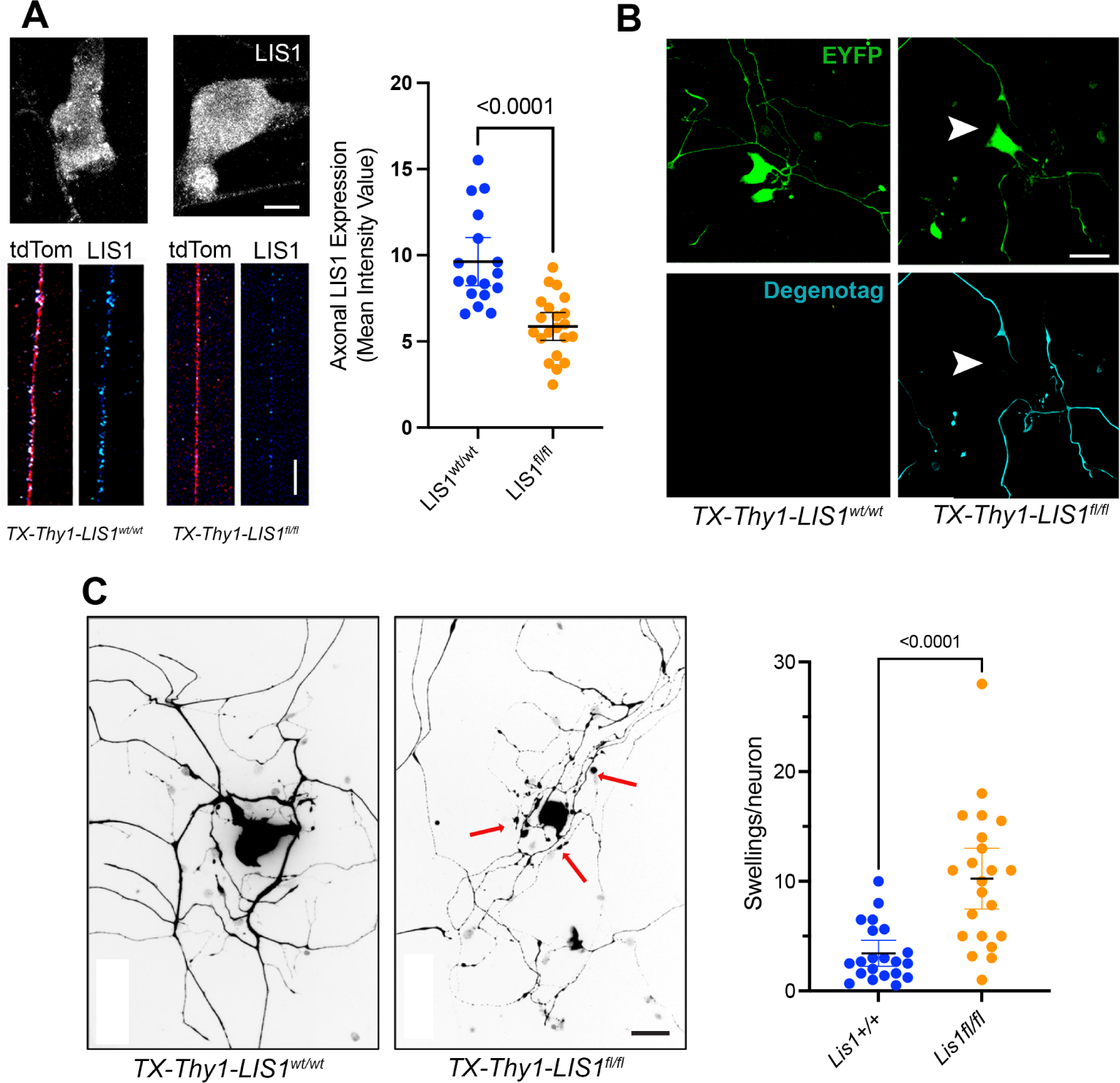
LIS1 depletion does not prevent axon growth in culture, but axons show signs of degeneration. **A)** Representative immunofluorescence images showing tdTomato-labeled axon (Red)) + LIS1 staining (Cell body in grey and axon in royal blue) in cultured DRG neurons from *TX-Thy1-LIS1^wt/wt^* and *TX- Thy1-LIS1^fl/fl^* mice received five doses of TX. Axons from *TX- Thy1-LIS1^fl/fl^* mice have greatly reduced LIS1 protein levels compared to control axons. Graph shows mean ± 95% CI for 3 biological replicates, 20 axons per group. Significance determined using a two-tailed Student’s t test. Scale bar = 20 µm. **B)** Cultured DRG neurons from *TX-Thy1-LIS1^wt/wt^*and *TX-Thy1-LIS1^fl/fl^* mice received five doses of TX and were stained with Degenotag (cyan) to visualize degenerating axons. Scale bar = 10 µm. **C, D)** In cultured DRG neurons, axons from *TX-Thy1-LIS1^fl/fl^* mice appeared more fragile and fragmented than controls and exhibited a significant inCreERT2 in swellings per neuron, consistent with axonal pathology. The graph shows the mean ± 95% CI of neurons from 3 mice per genotype. Significance was determined by two-tailed Student’s t test of measurements from 22 neurons per group. Scale bar= 10 µm.

Axonal transport of mitochondria is disrupted by LIS1 knockdown in cultured adult rat DRG neurons(Pandey et al., 2022). Transport of acidic organelles was also disrupted in rat DRG cultures, as well as in DRG neurons cultured from TX-*Act-LIS1^fl/fl^* mice (Pandey and Smith, 2011; Hines et al., 2018). In the current study we examined the steady state distribution of mitochondria and early endosomes using IF in fixed DRG cultures **(Fig. 8)**. We used and antibody against the mitochondrial flavoprotein AIF to label mitochondria. AIF intensity was stronger on growth cones from *TX-Thy1-LIS1^fl/fl^* mice pointing to decreased retrograde transport of these organelles out of growth cones **(Fig. 8A-C).** There was large variability in the distribution and shape of mitochondria control axon shafts **(Fig. 8D**). This variability was also observed in individual *TX-Thy1-LIS1^fl/fl^* axons, and the absolute number of mitochondria were not different between knockout and control axons **(Fig 8E)**. However, the average lengths of mitochondria per 100 µm axon segment was greater in *TX-Thy1-LIS1^fl/fl^* axons **(Fig. 8F)**, a phenomenon that was also documented after LIS1 knockdown in adult rat neurons(Pandey et al., 2022). The distribution of early endosomes labeled with EEA1 antibody was also changed by LIS1 depletion. Like mitochondria early endosome staining was more intense in *TX-Thy1-LIS1^fl/fl^* growth cones **(Fig. G, F).** While it was difficult to distinguish individual vesicles in growth cones, vesicles were easily observed in cell bodies **(left panel in G).** We did not observe a significant increase in EEA1 intensity in cell bodies, which suggest normal anterograde transport, although this was not tested directly. Taken together with our previous findings these data demonstrate that LIS1 depletion allows adult neurons to extend axons but compromises axonal transport and integrity.

**Figure 8:**
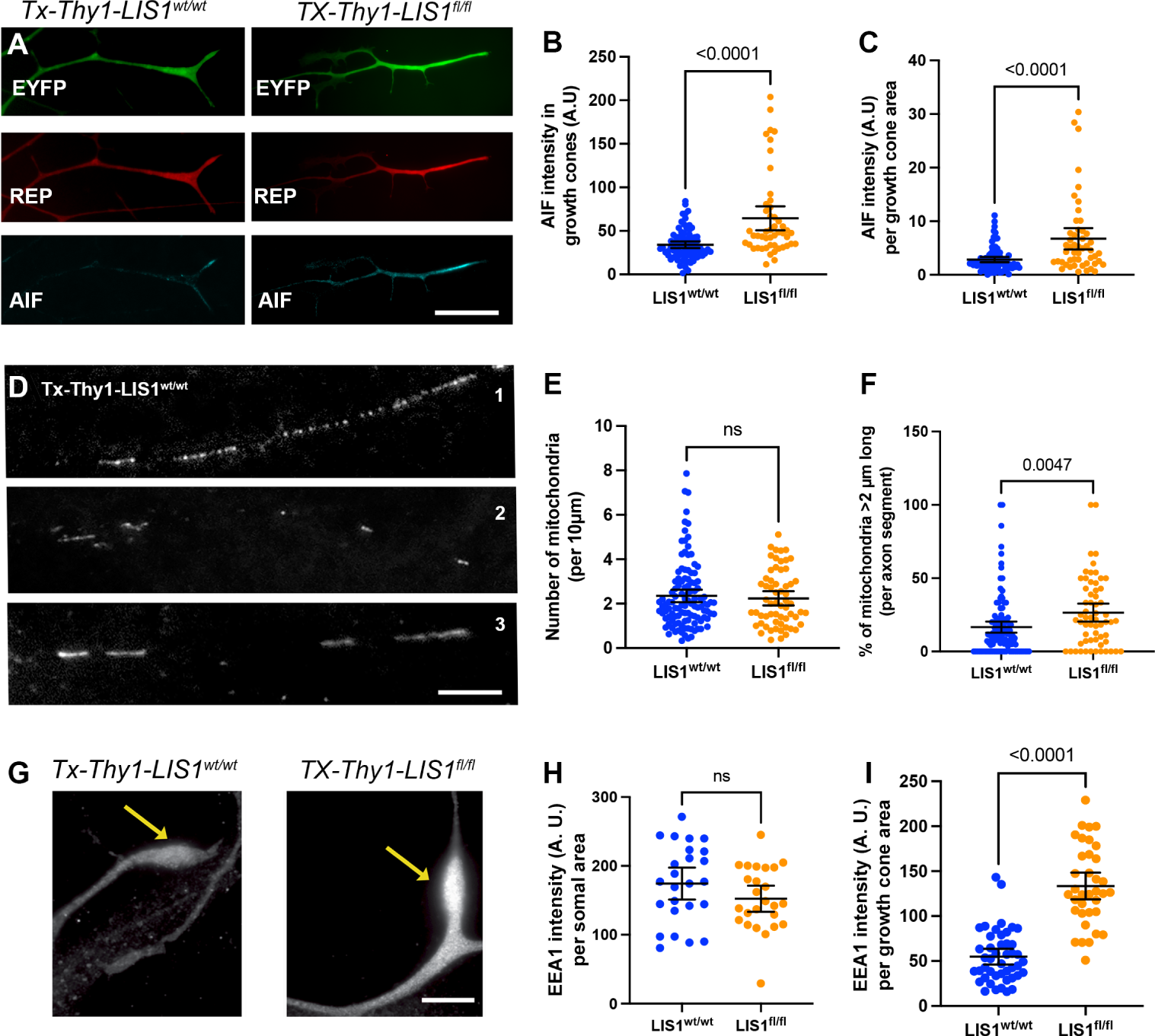
LIS1 depletion results in altered distribution of axonal mitochondria and early endosomes. **A**) Individual growth cones at ends of axons in DRG cultures from *TX-Thy1-LIS1^wt/wt^* and *TX-Thy1-LIS1^fl/fl^* mice (LIS1^wt/wt^ and LIS1^fl/fl^, respectively). Axons and growth cones from SLICK-H mice are positive for EGFP regardless of TX exposure. Both LIS1^wt/wt^ and LIS1^fl/fl^ express CreER and the tdTomato Cre activity reporter, so both show tdTomato expression when exposed to TX. LIS1^fl/fl^ growth cones show more staining for the mitochondrial marker, AIF. Average AIF intensity was measured for growth cones selected using a segmented line tool in ImageJ. Data is shown as the intensity per growth cone of 50 growth cones from 3 animals, **(B)** or as this intensity divided by the area of the region of interest **(C)**. Both showed more intense IF in LIS1^fl/fl^ growth cones. Significance determined by two-tailed Student’s t test; graphs show the mean ± 95% CI**. D)** Mitochondria in different axon shafts are not uniform in size or shape, and **E)** are not different in number in LIS1^wt/wt^ and LIS1^fl/fl^ axons. **F)** More axons in LIS1^fl/fl^ attain a length of >2 µm. Lengths were determined from plot profiles of segmented lines from 100 axon segments (100µm in length) from 3 different animals and include over 4000 individual mitochondria. The graph shows the % of mitochondria longer than 2 µm per axon segment. Significance determined by two-tailed Student’s t test; graphs show the mean ± 95%CI**. G)** Panels show growth cones (arrows) from LIS1^wt/wt^ and LIS1^fl/fl^ axons in the same culture labeled with EEA1. **H)** No significant difference was observed in EEA1 intensity in cells bodies. This was measured using circular ROIs of different sizes drawn to avoid the nucleus and cell cortex. The intensity measurements were divided by the area of the RO1, and the data shows intensity per area. I) EEA1 intensity in growth cones was determined as described above for mitochondria. Significance determined by two-tailed Student’s t test; graphs show the mean ± 95% CI. Scale bars = 10 µm (A), 5 µm (D, G).

## DISCUSSION

This study establishes that LIS1 plays a critical role in maintaining the structural and functional integrity of projection neurons in the adult nervous system, and that LIS1 also has important roles in adult astrocytes. We employed inducible knockout strategies to test this and found direct evidence that LIS1 depletion in projection neurons, but not in astrocytes, triggers rapid-onset axonal pathology, synaptic dysfunction, motor deficits, seizures, and premature lethality. These findings reveal an essential requirement for LIS1 specifically in adult mouse projection neurons and demonstrate that its role in regulating dynein activity extends well beyond development into the long-term preservation of neuronal health and maintenance of neuronal function. Dynein dysfunction contributes to axonal degeneration in human motor neuron diseases such as Charcot-Marie-Tooth type 20 (CMT20) and spinal muscular atrophy with lower extremity predominance (SMALED), and although this has not been demonstrated for LIS1, our studies may provide insight into potential therapeutic approaches to these disorders. Mutations that disrupt dynein processivity in mice also cause axonal degeneration (Ori-McKenney et al., 2010; Ori-McKenney and Vallee, 2011), so our current model is that the observed axonal degeneration phenotypes in adult *TX-Thy1-LIS1^fl/fl^* mice are caused by reduced dynein processivity. There is evidence that LIS1 supports a high force, high load-bearing state that would favor regulation of certain cargo classes, such as mitochondria and large vesicles (McKenney et al., 2010), so these organelles might be specifically relevant to the axonal degeneration phenotype observed.

A central observation of our study is that LIS1 depletion in projection neurons induces axonal swellings, fragmentation and accumulation of neurofilament degradation products characteristic of the Wallerian degeneration that occurs in distal axons after axotomy. These degenerative changes were not restricted to distal axons, as would be typical in a “dying-back” model of neurodegeneration, but instead occurred along the entire axonal length. This suggests that LIS1 loss disrupts dynein-mediated axonal transport globally, impairing the ability of projection neurons to maintain axonal homeostasis. SARM1 (Sterile alpha and TIR motif containing 1) is an NADase implicated in Wallerian degeneration after nerve injury (Gilley et al., 2015; Sasaki et al., 2016; Figley et al., 2021). Its activity is stimulated by the failed delivery of NMNAT2 (nicotinamide mononucleotide adenylyltransferase 2) into severed axons. NMNAT2 normally converts NMN (nicotinamide mononucleotide) and ATP to NAD+, whose presence is critical for metabolic homeostasis, driving both mitochondrial function and on-board vesicle glycolysis during axonal transport. Axon injury prevents transport of Golgi-derived vesicles containing NMNAT2 into severed distal axons leading to a buildup of NMN. NMN stimulates SARM1’s NADase activity, leading to NAD+ depletion, ATP loss, and reduced mitochondrial motility and respiration (Ko et al., 2021). This then causes calcium influx, membrane permeability, calpain activation and axonal fragmentation. It seems unlikely that LIS1 would inhibit SARM1 directly. We’ve previously demonstrated that in cultured DRG neurons from adult rats LIS1 knockdown also reduced anterograde transport of both acidic organelles and mitochondria. However, we did not detect a decrease in anterograde transport of acidic organelles in DRG axons in cultures from TX-Act-LIS1^fl/fl^ – in fact anterograde transport was modestly but significantly increased (unpublished data). Future studies will be required to determine if canonical SARM1 pathways are involved in the degeneration of LIS1 depleted axons, or whether the pathology arises from defective retrograde transport of spent mitochondria and other organelles without activation of SARM1. Indeed, we have observed an increase in intensity of AIF and EEA1 IF in growth cones of cultured neurons, indicating that potentially spent organelles are not being adequately removed.

The graded effects of tamoxifen dosing allowed us to measure changes that occurred in axons over time. Mice receiving five consecutive tamoxifen injections displayed rapid, widespread and severe axonal degeneration. Animals needed to be euthanized by 10 days after the first injection, whereas mice exposed to fewer injections exhibited milder phenotypes and prolonged survival. This dose-dependent effect corresponds to the proportion of projection neurons undergoing recombination. Because of the complex cytoarchitecture and many cell types in the adult brain our finding underscores the importance of using a CreERT2 activity reporter to reveal which cells have undergone CreERT2 mediated recombination. The slower decline allowed us to determine that the pathological changes in individual axons became progressively worse with time.

Another important aspect of this work is the comparison of the phenotypes caused by LIS1 depletion in projection neurons and astrocytes. While neuron-specific LIS1 knockout led to severe neurological decline and death, astrocyte-specific LIS1 knockout did not produce any overt motor deficits or reduction in lifespan. The elevated GFAP expression observed in LIS1 depleted astrocytes is intriguing and demonstrates the functional significance of LIS1 in this cell type in adults. GFAP upregulation is a hallmark of “reactive astrogliosis” which occurs in response to stress, injury or inflammation in the CNS. By definition, reactive astrogliosis is secondary to an extrinsic signal (Escartin et al., 2021). We did not intentionally provide an external signal so this LIS1 depletion response may not be classic reactive astrocytosis. However, if LIS1 loss in astrocytes perturbs their homeostatic roles in the brain, neurons may in fact develop pathologies and signal back to the astrocytes. GFAP upregulation is thought to contribute to astrocyte hypertrophy and proliferation in response to stress. It is possible that LIS1 (directly or indirectly through dynein) impacts GFAP transcription, translation or stability, but we have no direct evidence for this. It will be interesting to determine if LIS1 expression is downregulated in astrocytes when astrogliosis is triggered by a different perturbation. If so, LIS1 downregulation may be an important event in astrogliosis, either serving as a protective strategy or driving inflammation.

An alternative scenario is that LIS1 depletion is sufficient to trigger reactive astrogliosis, indirectly leading to GFAP upregulation. Further studies will be needed to determine if other markers of astrogliosis are present and if morphological changes accompany the increased GFAP expression. Our data indicate that LIS1 depletion in astrocytes impacts dynein distribution and MT trafficking. It will be interesting to determine if astrocytic functions like phagocytosis and synaptic pruning are impacted by LIS1 depletion, and if astrocytic LIS1 depletion causes any behavioral changes in mice. Disruption of dynein-dependent processes like vesicular trafficking and protein transport might also be involved in reactive astrogliosis. For example, defective trafficking of junctional proteins could impact BBB integrity, or altered trafficking of mitochondria within astrocyte processes could impact metabolism and trigger a reactive phenotype.

Finally, perturbation of an important LIS1/dynein-dependent process in astrocytes could have a pathological effect on neighboring neurons, which in turn could trigger astrogliosis. We have not detected gross anatomical abnormalities in these mice, but as the mice survive long term, any axonal or synaptic pathology would likely be more modest than observed after LIS1 depletion in projection neurons. Nonetheless it will be interesting to look for more subtle or localized impacts on axons and synapses.

Taken together, our findings suggest that LIS1 sustains neuronal viability in adulthood by supporting dynein-mediated transport and possibly preventing activation of intrinsic degeneration pathways. Disruption of LIS1 and dynein dependent processes at the AIS might contribute to the phenotypes observed in TX-Thy1-LIS1^fl/fl^ mice. LIS1 and dynein are important for the rerouting of misrouted dendritic components out of axons and also contribute to the polarity sorting of MTs entering axons (Kuijpers et al., 2016; Klinman et al., 2017; Jenkins and Bender, 2024).

Although less well studied, LIS1 also functions in dendrites, at least during development (Kawabata et al., 2012; Sudarov et al., 2013). However, given the rapid onset of axonal degeneration, we consider disrupted axonal transport to be the most likely event contributing to the neurological phenotypes. Regardless, loss of LIS1 appears to tip the balance toward axonal self-destruction, with catastrophic consequences for neuronal and organismal survival.

In conclusion, our findings establish that LIS1 is a vital regulator of neuronal survival well into adulthood. Its depletion leads to rapid axonal degeneration and neurological decline through mechanisms likely involving disrupted dynein activity. This work extends the functional scope of LIS1 beyond development, defining it as a lifelong guardian of axonal integrity in long-range projection neurons. This also provides new avenues for investigating therapeutic interventions aimed at preserving axonal health in neurodevelopmental and neurodegenerative disorders.

## Acknowledgments

We thank Dr. Jeffrey Twiss for comments, suggestions and reagents.

---

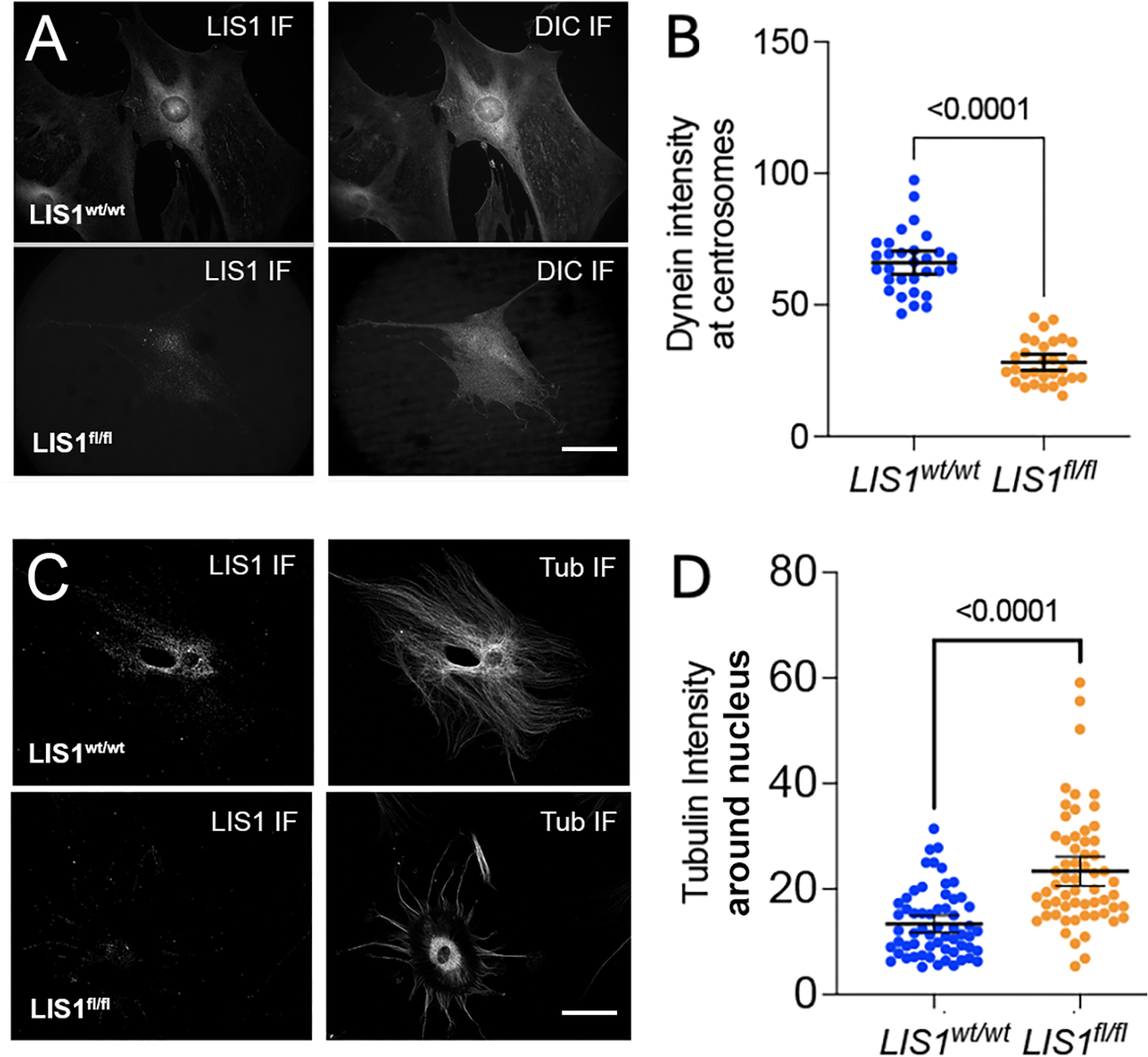

**Movie S1:** TX-Thy1-Lis1wt/wt do not have neurological phenotypes on day 10 after the first of 5 tamoxifen injections.

**Movie S2:** TX-Thy1-Lis1wt/wt have neurological phenotypes on day 10 after the first of 5 tamoxifen injections. These include shivering, altered gait, Straub tail, which is typical of seizure activity.

